# Regulation of thickness of actomyosin cortex in well-spread cells by contractility and spread area

**DOI:** 10.1101/205138

**Authors:** Rinku Kumar, Bidisha Sinha

**Affiliations:** Department of Biological Sciences, Indian Institute of Science Education and Research (IISER) Kolkata, Mohanpur 741246, West Bengal, India

**Author notes:** Author for correspondence: Bidisha Sinha.

## Abstract

The contractile cortical actomyosin cytoskeleton (or cortex) in interphase cells confers rigidity to cells, but also lead to shape dynamics. Regulation of its thickness, although well studied in rounded cells, is less explored in well-spread cells. In this paper, we quantify the variations in thickness and study the contribution of actin polymerization, myosin II activity and spread area of cells. We report an increase in cortex thickness and its variations on disrupting actin network by actin depolymerizing agents or reducing contractility by inhibiting motor activity of myosin II. On spread area reduction by substrate micropatterning, we find reduced cell volume and increased mean & variability of thickness. To validate, we follow cells through de-adhesion with EDTA. The thickness of cortex increases (and oscillates) while the volume of cells reduces with 5-15 mins timescales. Moreover, total internal reflection fluorescence (TIRF) imaging reveals stress fibre dissolution and events of their buckling along with a growing population of micron-sized mobile filaments. We believe that the cytoskeleton responds to the loss of adhesion by contracting and fragmenting, hence leading to cortex thickening. Limiting volume reduction does not suppress cortex thickening on de-adhesion, suggesting that decreased traction stress may be primarily responsible for the cortex thickening.

## Introduction

The cellular cortex is a thin layer of actin filaments under the plasma membrane crosslinked by myosin II and actin binding proteins (1, 2). Its architecture is distinct from other actin-based structures that may also exist at the membrane such as, stress fibres, filopodia or lamellipodia (2). Anchored to the membrane through Ezrin, Radixin, and Moesin proteins (3), the actin filaments of the cortex are disordered but lie mostly in the plane of the membrane. The network is contractile due to the motor activity of myosin II and may have a tension of ∼ 500 pN/μm in fibroblasts (1). The thickness of the cortex, ∼300-1000 nm (1, 4), is established by the polymerization/depolymerization of actin (5). However, it can be dynamically altered due to the turnover of actin filaments and associated crosslinkers such as myosin II, α-actinin and formins in interphase cells (6) as well as length regulators like CFL1, CAPZB and DIAPH1 in mitotic cells (7). Although the cortex provides rigidity to elicit protection to the cell (3, 8), it can hinder the trafficking of vesicles from cytoplasm to plasma membrane and *vice versa* (9, 10). An asymmetrically contractile cortex, moreover, can potentially generate shape oscillations (11) and mechanical heterogeneity may favour fragmentation (12). The regulation of the cortex – at a proper thickness and spatial homogeneity is therefore crucial for the working of the cell.

Thickness regulation has been mostly studied in geometrically round cells (7) but not extensively in well-spread cells. In round cells, cortex thickness is inversely correlated with cortex tension during progression through mitosis and regulated by the actin length-regulators and myosin II motor activity. In comparison to round cells, well-spread cells experience larger traction stresses from the substrate. As adhesion progresses, the round geometry of cell is replaced by a flattened geometry and substrate forces become dominant over the rounding effect of Laplace pressure. Focal adhesions are well formed and stress fibres are more prominent in these cells (13). Since myosin II binding to actin favours higher tension (14), increased traction stress may alter thickness regulation in well-spread cells. Therefore, the regulation of cortex thickness needs to studied keeping in mind the strong forces from the extra cellular matrix. In this study, well-spread cells are used to understand the regulation of cortex structure, thickness and its heterogeneity.

Interphase CHO cells present the right cortical morphology and pharmacological agents help in altering the actomyosin activity at the cortex. Further, using micropatterning and EDTA, cells are transited from suspended to adhered states (with restricted spread area) and from adhered to de-adhered states respectively. Using epi-fluorescence, we assess the cortex thickness and using TIRF microscopy, we quantitate changes in the architecture of the cortical network.

## Materials and methods

### Cell culture, actin labelling and image acquisition

All experiments were conducted using CHO cells cultured in Dulbecco’s Modified Eagle’s Medium (Gibco, Life Technologies, USA) supplemented with 10 % Fetal Bovine Serum (Gibco). Cells were allowed to grow for 24 hrs before experiments unless otherwise specified. Fixation was performed using 4% paraformaldehyde (PFA) (Sigma) for 15 mins at 37°C followed washing thrice with PBS (Sigma). F-actin labelling was performed by first permeabilizing cells by 0.2 % Triton-X 100 (Sigma) for 5 mins, washing thrice with PBS wash and incubating with 33 nM Rhodamine-phalloidin (Molecular Probes, Life Technologies) for 45 mins in the dark. Coverslips were either fixed on slides with Mowiol (Sigma) for epifluorescence imaging or cells on glass-bottomed petri dishes were fixed and immersed in PBS for TIRF imaging. Images in all experiments were acquired on an inverted microscope (IX81, Olympus Corporation, Japan) using 100X TIRF objective (N.A 1.49) and a CMOS camera (ORCA-Flash 4.0, Hamamatsu Photonics, Japan) with a 1 pixel= 65 nm either in the epi-fluorescence or TIRF mode using DPSS lasers of 488 nm and 561 nm wavelengths. For z-stack acquisition, 0.5 μm z-step was used and for TIRF, penetration depth was kept at 70 nm. For live imaging, cells were maintained at 37°C throughout (Okolab, Italy).

### Plasmids and transfection

PDEST/Life-Act-mCherry-N1 (a gift from Rowin Shaw) (Addgene, plasmid # 40908) was used for actin labeling in live cells (15). pEGFP-C1 and pDsRed-Express2 (Clonetech Laboratories, USA) were used for cytosol labelling. CHO cells were plated on customized 35 mm glass bottom imaging dishes for at least 16 hr before transfection. All transfections were performed using 0.5 - 0.75 μg plasmid DNA by lipofection (Lipofectamine 3000, Life Technologies) according to manufacturer’s instructions.

### Perturbations using drugs

To perturb actin, cells were treated with 5 μM cytochalasin D (Cyto D) (Sigma) (16, 17), 5 μM latrunculin B (Lat B) (Sigma) (17) and 5 μM blebbestatin (Blebb) (Sigma) (18) for 30 mins before fixation and staining. For ATP depletion (ATP dep) cells were incubated with mixture of 10 mM 2-Deoxy-D-glucose and 10 mM NaN_3_ for 30 mins in glucose free M1 medium (19). 50 nM Jasplakinolide (Jaspla) (Life Technologies) (20) and 80 μM Dynasore (Dyna) (Sigma) (21) were used to prevent actin dynamics and Dynamin dependent endocytosis respectively.

### Surface Micropatterning And Cell Seeding

Glass coverslips were etched with 19:1 mixture of ethanol and acetic acid for 30 mins followed by washing twice and incubating for 15 mins with ethanol. Air dried coverslips were exposed with deep UV in the UV-Ozone cleaner for 5 mins (Jelight Company, USA). Clean coverslips were incubated with 0.2 mg/ml PLL-g-PEG (SuSos, Switzerland) (prepared in 10 mM HEPES (Sigma), pH 8.5) for 2 hrs inside a moisture box. Photo-masks (JD Photo Data, UK) already cleaned in the UV-Ozone cleaner for 5 mins were used for patterning the PEG coated coverslips for 10 mins. Coverslips with water drops were placed on the photomask and excess water soaked off to aid adherence. Patterned coverslips were removed from photomask by floating them off using water. Finally, patterned coverslips were incubated with 20 μl of 25 μg/μl fibronectin (Sigma) (prepared in 100 mM NaHCO_3_, pH 8.6) for 45 mins inside a moisture chamber. Patterned coverslips were washed twice with PBS before seeding cells. CHO cells were detached from culture using 0.02 % EDTA (Cal Biochem). The cell suspension was centrifuged to discard EDTA. 2 × 10^5^ cells were seeded per 25 mm coverslip and unattached cells were removed 30 mins after plating using equilibrated media. The micropatterning protocol was adapted from (22, 23).

### Cell de-adhesion

For measuring the evolution of cortex thickness and volume of single live cells through deadhesion, LifeAct-mCherry transfected CHO cells were treated with EDTA solution and followed. EDTA was prepared in PBS by adding 1 mM EDTA along with 20 mM glucose (Sigma). Imaging media (DMEM + 10% FBS) was replaced by EDTA solution and z-stacks were captured either before and after 1 hr of EDTA addition or every 5 mins after addition of EDTA. For understanding the effect of Dyna and Jaspla on de-adhesion, same cells were assayed in three phases - without drugs, with drugs in media and with drugs in EDTA solution at time points 0, 30 and 90 mins. For following only volume alterations on EDTA treatment, CHO cells, transiently transfected with pEGFP-C1 or pDsRed-Express2, were treated with EDTA and a particular plane (at ∼ 2-2.5 μm above the basal plane) was imaged every 2.5 mins.

### Image analysis and quantification

Image analysis was performed using softwares - Image J/Fiji (https://imagej.net/Fiji) and MATLAB (MathWorks). For calculating the full width at half maximum (FWHM) of cortex, the analysis was performed in two parts - linearization of the cell edges using Fiji and extraction of intensity line-scans and fitting them for FWHM calculation using MATLAB. Lines of 2 μm (31 pixels) length following cell cortex were drawn, fitted with non-uniform cubic spline and straightened. Pixels normal to the line (5 μm (77 pixels) across) were extracted for every point on the line and together represented the straightened cortex. Fluorescence line-scans perpendicular to the membrane were fitted using 3- and 4-term Gaussian function and the better fit (higher R^2^) used for further analysis. FWHM of the Gaussian whose peak was located closest to the centre of the line was used as a measure of thickness at that cross-section. FWHM averaged across ROIs or cells was termed the average thickness and the standard deviation (SD) termed the variability of thickness. For quantification of actin intensity, images of actin labelled cells were acquired using same concentration of Rhodamine-phalloidin and same acquisition parameters. Peak intensity values of line-scans were measured after background subtraction for both fixed and live cells. For relative changes in actin intensity in live cells, same cells were followed and same acquisition settings were used before and after EDTA administration. For spread area, volume and surface area quantification, stacks of actin labelled cells were captured from bottom to top using 0.5 μm step size. Manual ROIs along the boundary of each stack were drawn. Total projected area at the bottom plane were considered as a spread area of cells. Volume of each cells were computed by calculating the summation of the cell-areas obtained for each image of the stack and multiplying it by the z-step-size. Similarly, surface area was calculated by summing up the cell-perimeters obtained for each image of the stack and multiplying by z-step-size. Average parameters termed energy, orientation and coherence reflect the total intensity, the orientation and coherence in pixel intensity distributions (24) that were calculated for each frame of image sequences using Orientation J plugin of Image J. SD of orientation for 5 consecutive frames (using sliding window algorithm) was calculated and termed SD(Orientation). Single filament analysis was performed by first normalizing images (0.3% saturated pixels) in Fiji and then detecting objects by background subtraction followed by local thresholding using Otsu thresholding algorithm using MATLAB.

### Endocytosis quantification

Petri dishes with CHO cells were kept on ice for 10 mins before administration of 5 μM FM 1-43 FX (Molecular Probes) resuspended in PBS. Cells were incubated for 1 hr and fixed for 15 mins using 3.7 % PFA. For control, PBS supplemented with glucose (20 mM) was used as a medium and for de-adhesion, EDTA (1 mM in PBS supplemented with glucose). Excess free FM 1-43 FX was washed with PBS before fixation. Total amount of endocytosed FM 1-43 FX was calculated by integrating the fluorescence intensity of each cell over whole stacks captured with 0.5 μm z-step-size.

## Results

### Cortex thickness variations across single cells and among a population are similar

We measure cortex thickness in CHO cells by computing FWHM of the F-actin close to the membrane at cell edges (Fig. 1 A). The well-spread interphase cells are chosen and analysis performed (Fig. 1 B, C) at regions that lack specialized structures like lamellipodia or filopodia. We calculate the thickness at a region of interest (ROI) (eg. rectangle in Fig. 1 A) by straightening the cortex (Fig. 1 B, *top*, *middle*) and fitting line-scans normal to the membrane at each pixel along the cortex (Fig. 1 B, *bottom*). The FWHM of the Gaussian centred at the peak of the profile is determined and used as a measure of the cortex thickness. Across a typical cell, multiple ROIs are considered for analysis (Fig. 1 C). Thickness values are obtained from individual line-scans for all ROIs from a single cell (Fig. 1D *left*) and compared with values of average thickness per ROI obtained from 30 different (Fig. 1D *right*). The average thickness for single ROIs in a particular cell (Fig. 1D *left*, dotted line) is observed to be ∼ 382 ± 57 nm (error denotes SD). In comparison, thickness averaged over different cells was calculated be ∼ 424 ± 62 nm (Fig. 1D, *right*, dotted line). Hence, cortex thickness and variability (SD) across single cells is similar to those among a population. We confirm that this measurement is independent of the signal-to-noise ratio of the images (Fig. S1 in the Supporting Material). It’s important to note that the FWHM may overestimate the thickness due to the wide point spread function (PSF) of the imaging system. Hence the numbers need to be compared in relative terms.

**FIGURE 1.**
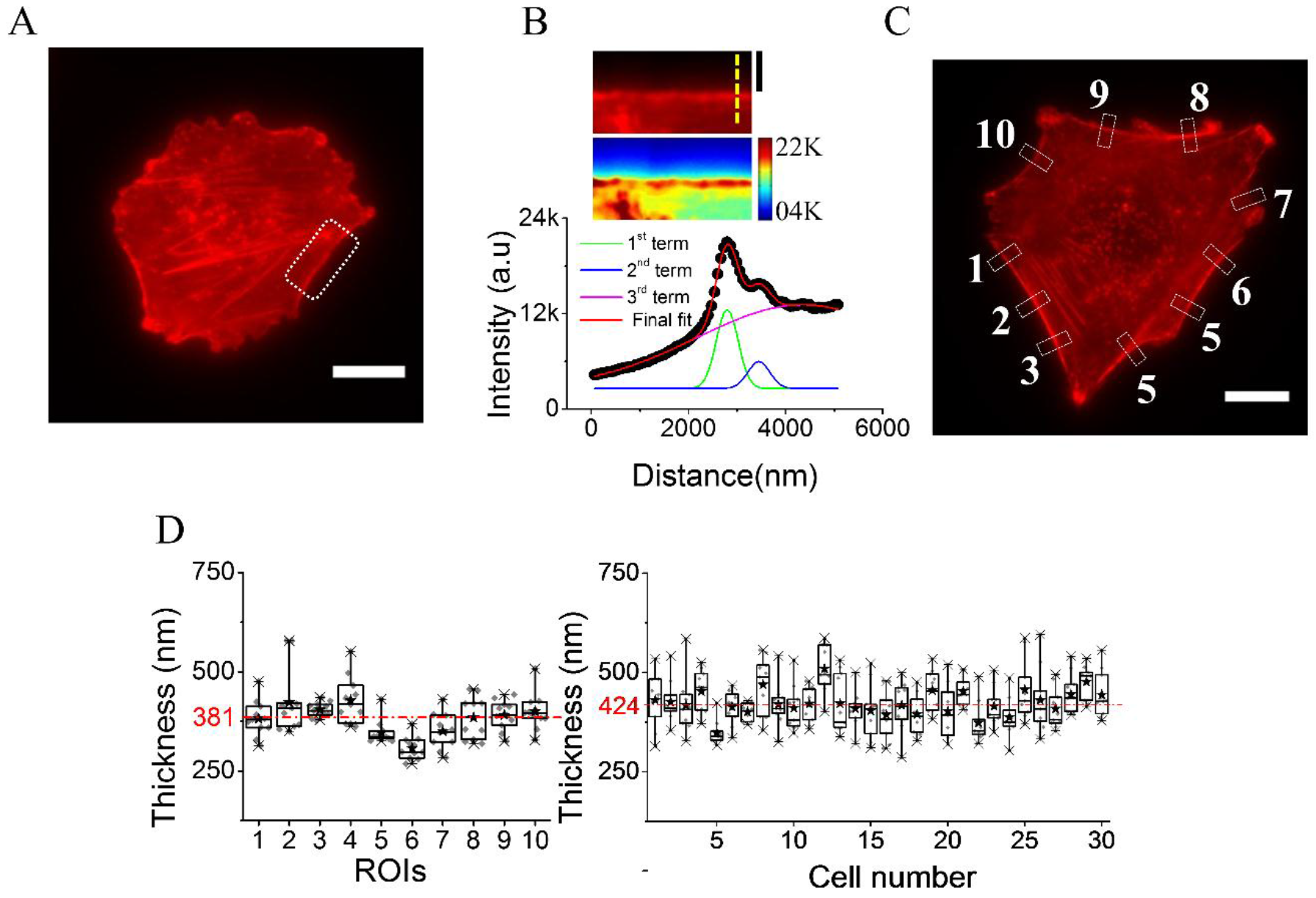
Measurement of cortex thickness. (A) Representative image of a CHO cell stained with Rhodamine-phalloidin. Scale bar, 10 μm. (B) Top: Zoomed-in view of dotted ROI in (A) after linearization of the cortex. Scale bar, 2 μm. Middle: Color coded image of intensity. Bottom: Intensity line-scan of the linearized cortex (at yellow dashed line of (B)) and fit to a 3-term Gaussian. (C) Image of CHO cell stained with Rhodamine-phalloidin with ten random ROIs (2 μm along the cortex and 5 μm across) marked out. Scale bar is 10 μm. (D) Left: Box plot of cortex thickness derived from line-scans for the 10 ROIs (as marked in C) and Right: Box plot of average cortex thickness of single ROIs for 30 cells (each point in box plot represents mean of at least 11 line-scans (2 μm) along the cortex). Error bars represent SD.

To understand the mechanisms involved in maintaining the cortex thickness, the next part of our study focusses on the relative changes observed in cortex thickness on drug-based biochemical perturbations.

### Cortex thickness increases on actin depolymerization and reduced myosin II activity

We treat cells with Cyto D or Lat B to enhance depolymerization and to additionally prevent polymerization of actin respectively. We also employ ATP depletion to block all ATP dependent activities and Blebb to block myosin II motor activity (18). Cyto D or Lat B treatments result in reduced actin intensity at the cortex with visible heterogeneities due to removal of actin from parts of the cortex and formation of clumps at other parts (Fig. 2 A). This is visible in the straightened image of Cyto D or Lat B treated cells (Fig. 2 B) and consistent with results proposing myosin II motor activity contracting the weakened cortex into clumps (25). Time-lapse imaging of LifeAct-mCherry expressing cells show the rupturing and clumping of the cortex (Movies S1 & S2 in the Supporting Material). In contrast, ATP depletion and Blebb treatment enhance the F-actin intensity at the cortex. Comparison of typical line-scans across the cortex (Fig. 2 C) show reduced peak intensity for Cyto D or Lat B treatments while increased peak intensity for ATP depletion and Blebb treatments (Fig. 2 D). The role of activity in maintaining a reduced cortical intensity of actin is, thus, implicated. FWHM measurements show that the average thickness increases on all the treatments (Fig. 2 E). The distributions display that the variability of thickness also increases on all these treatments (Fig. 2 F). Hence, weakening of the network both by reducing actin polymerization or blocking myosin II results in increased heterogeneity of cortex thickness across single cells. This is consistent with myosin-II’s role in creating lateral coherence in the cortical structure (26). Analysing time-lapse images of a cell undergoing cortex rupture by Cyto D, we find that the actin concentration profile spreads out while the thickness increases with time (Fig. S2 A-C in the Supporting Material) while the cortex ruptures at 0.5-1.5 μm/min (Fig. S2 D-E). Rupturing is sometimes preceded by straightening of the cell-edge – indicating build-up of tension - accompanied by a transient reduction in cortex thickness (Fig. S2 F-H).

**FIGURE 2.**
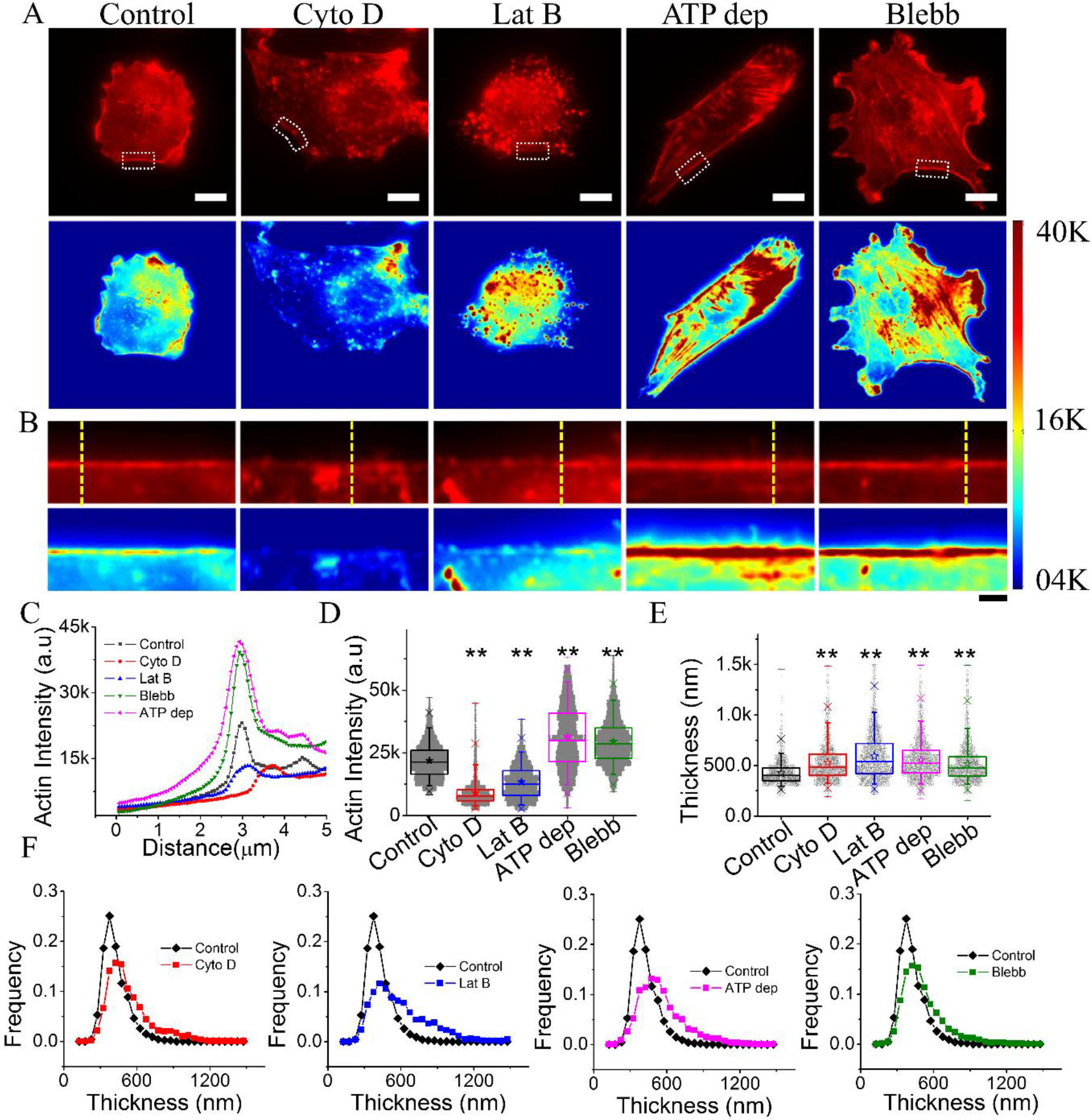
Effect of actin and/or myosin perturbation on cortex. (A) Representative images of Control, Cyto D, Lat B, Blebb treated and ATP depleted CHO cells stained with Rhodamine-phalloidin (top) and corresponding color coded image (bottom). Scale bar, 10 μm. (B) Linearized and zoomed-in view of the cortex (at indicated ROIs in (A)) and their color coded representation. Scale bar, 2 μm. (C) Line-scans across cortex (at yellow ROIs marked out in (B)) for control cells and cells under different indicated treatments. (D) Scatter box plot of peak cortex actin intensity line-scan after background subtraction, measured for 30 cells for each set with 10 ROIs per cell and 31 line-scans (2 μm) per ROI (total 9887, 9729, 9858, 9659 and 9903 line-scans for Control, Cyto D, Lat B, ATP dep and Blebb respectively). (E) Scatter box plot of thickness measured for 30 cells for each set with 10 ROIs per cell and11 line-scans (2 μm) per ROI (total 2536, 2051, 3363, 2466 and 2094 line-scans for Control, Cyto D, Lat B, ATP dep and Blebb respectively). (F) Distribution of thickness of Control, Cyto D, Lat B, ATP dep and Blebb treated cells. One-way ANOVA using Bonferroni post hoc method was performed with p cut off value < 0.05 and p<0.001 (denotes * and **respectively). Error bars represents SD.

### Higher cortex thickness when spreading is restricted to smaller area

Stress fibres link distal focal adhesions with contractile actomyosin structures and play a crucial role in imparting traction stress. Spread area alterations has been reported to result in different stress fibre density and also altered levels of maximum traction stress (27) and strain energy (28). To understand how the cortex responds to changes in cell’s spread area, we use micropatterns of varying adhesion areas (RecA: 5 μm × 10 μm; RecB:15 μm × 30 μm, Fig. 3 A *inset*) and monitor cortex thickness at 1 hr and 24 hrs post plating (Fig. 3 A). The normalized line-scans (Fig. 3 B) show increased widths for cells on micropatterns in comparison to the well-spread control cells. While the spread areas of the cells for both patterns are smaller than the control (Fig. 3 C), 3D volume rendering (Fig. 3 A) and quantification (Fig. 3 D) show that they have significantly reduced volumes and increased cortex thickness (Fig. 3 E) than control cells on glass. However, there is a relative increase in spread area for cells imaged 24 hrs post plating compared those imaged at 1 hr post plating for micropatterns and a concomitant increase in volume and decrease in thickness. Correlating the thickness for each cell against its volume and spread area (Fig. 3 F *left, centre*) a power law dependence of −0.248 (adjusted R^2^ = 0.43) defines the overall dependence of cortex thickness on cell volume when all the conditions are considered together. For individual conditions only the control cells have a −0.145 (adjusted R^2^ = 0.114) as the exponent while others appear uncorrelated with exponents ranging from −0.02 to 0.02 (Adjusted R^2^ ∼ −0.02 to −0.01). We find that thickness shows similar dependence with spread area with the overall exponent ∼−0.1 (Adjusted R^2^ = 0.49) while individual conditions are uncorrelated with exponents ranging from ∼ −0.08 to −0.1 (Adjusted R^2^ ∼ −0.03 to 0.05). The spread area and volume (Fig. 3 F, *right*) are also correlated with a power law exponent of 0.317 (adjusted R^2^ = 0.65). Except for well spread cells (exponent: 0.68, adjusted R^2^ = 0.45), in all other conditions spread area is uncorrelated with volume (exponent: −0.08 to 0.06, adjusted R^2^ = −0.02 to 0.008).

The distribution of thickness for cells on micropatterns display increased widths than control cells (Fig. 3 G). Tail regions are overrepresented in cells imaged 1 hr post plating than those imaged at 24 hrs post plating. The increased width/SD may indicate a loss of lateral coherence of the cortex which however partly recovers after 24 hrs. implying that the cortex thickness achieved in a cell, that goes from a suspended to an adhered state, is closely dependent on the final adhesion constraints. We next probe the response when the sequence of events is inverted - ie, cells are taken from their well spread geometry to a deadhered state.

**FIGURE 3.**
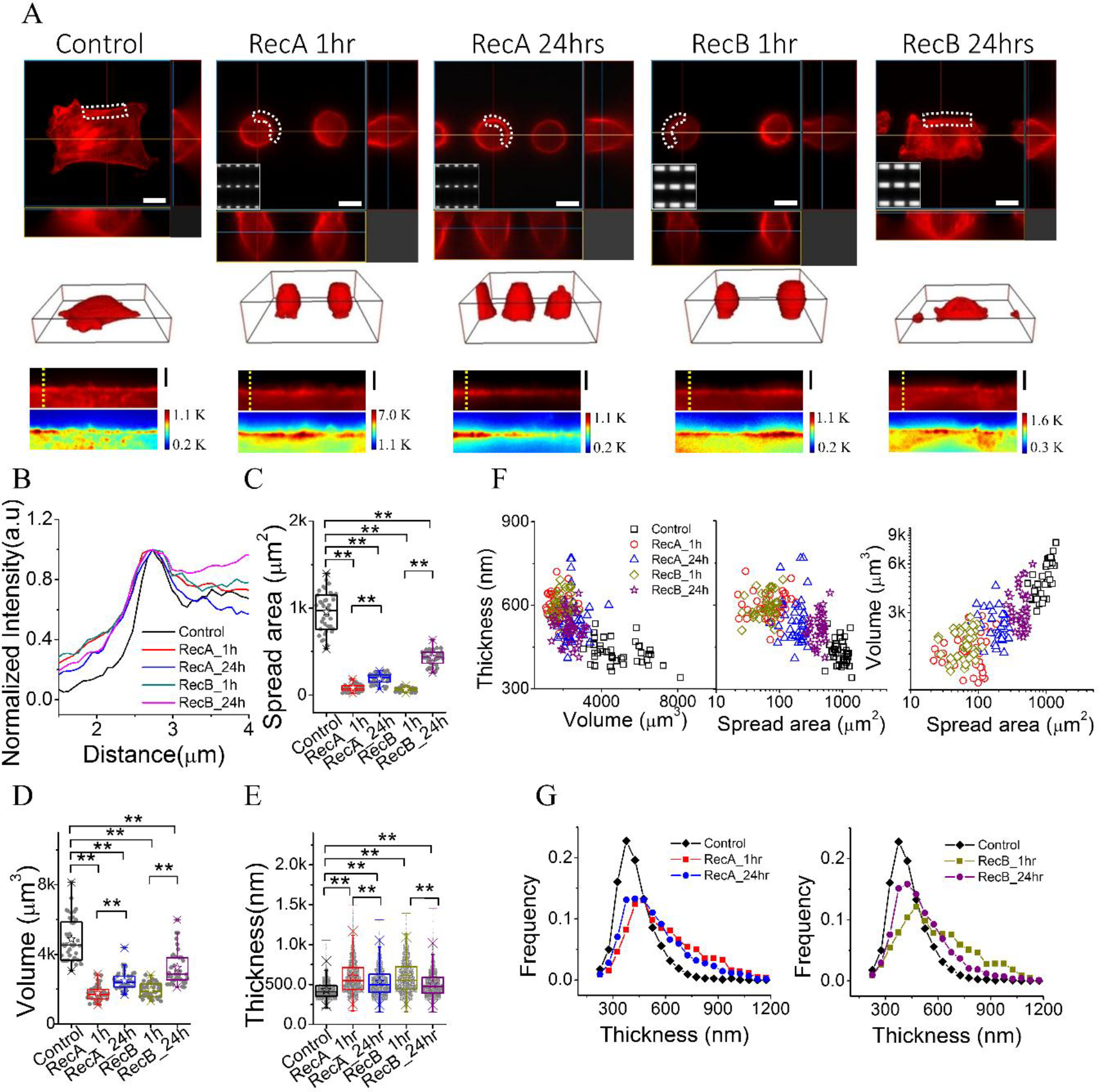
Cortex thickness increases for smaller spread area. (A) Top: Slice view (xy, xz and yz planes) of fixed and Rhodamine-phalloidin stained CHO cells grown on glass and micropatterns RecA and RecB for 1hr and 24h hrs as indicated. (Insets: Brightfield images of photomasks with RecA (10 μm x 5 μm) and RecB (30 μm x 10 μm) patterns). Scale bars, 10 μm. Middle: 3D rendered volume view. Bottom: Linearized and color-coded view of the cortex of cell region depicted as dotted ROI. Scale bars, 2 μm. (B) Comparing normalized intensity profile obtained from corresponding yellow dotted ROI in (A). (C) Spread area, (D) Volume and (E) Cortex thickness of cells. (F) Scatter plot of volume with cortex thickness (left), spread area with cortex thickness (middle) and spread area with volume (right). (G) Distributions of cortex thickness of cells as indicated. Error bars represent SD. Wilcoxon based Mann-Whitney U-test were performed for finding statistical significant difference in all cases except thickness measurement (one-way ANOVA using Bonferroni post hoc method) with p cut off value < 0.05 and p<0.001 denotes * and **respectively.

### Cortex thickness increase on de-adhesion

We perform experiments in live cells in which the cortex is labelled by Lifeact-mCherry and the cortex thickness, spread area and volume are measured on de-adhesion induced by incubation with EDTA. Multiple cells are either imaged at a time window post EDTA addition or single cells are followed in time. 3D rendered images of z-stacks capture the rounding of cells also evidenced as the enhancement of projected z-height of cells before and post 60 mins of de-adhesion (Fig. 4 A). The spread area of cells reduces post de-adhesion along with reduction in cell volume (Fig. 4 B, *top*). The intensity of F-actin at the cortex shows a slight but significant increase but the cortex thickness clearly increases on de-adhesion (Fig. 4 B, *bottom*). We again find no strong correlation between cortex thickness and either volume or spread area in the separate conditions although combining both suggest thickness to be inversely proportional to both. (Fig. 4 C, *left, centre*). The spread area and volume show a direct correlation as found earlier (Fig. 4 C, *right*). The distribution of thickness in round cells widen with greater contribution from higher values of thickness (Fig. 4 D). To explore the mechanism of thickness increase, we next follow the deadhesion dynamics by imaging particular cells through de-adhesion (Fig. 4 E).

We observe that the spread area reduces exponentially with timescales ∼ 5-20 mins (Fig. 4 E *left*).

**FIGURE 4.**
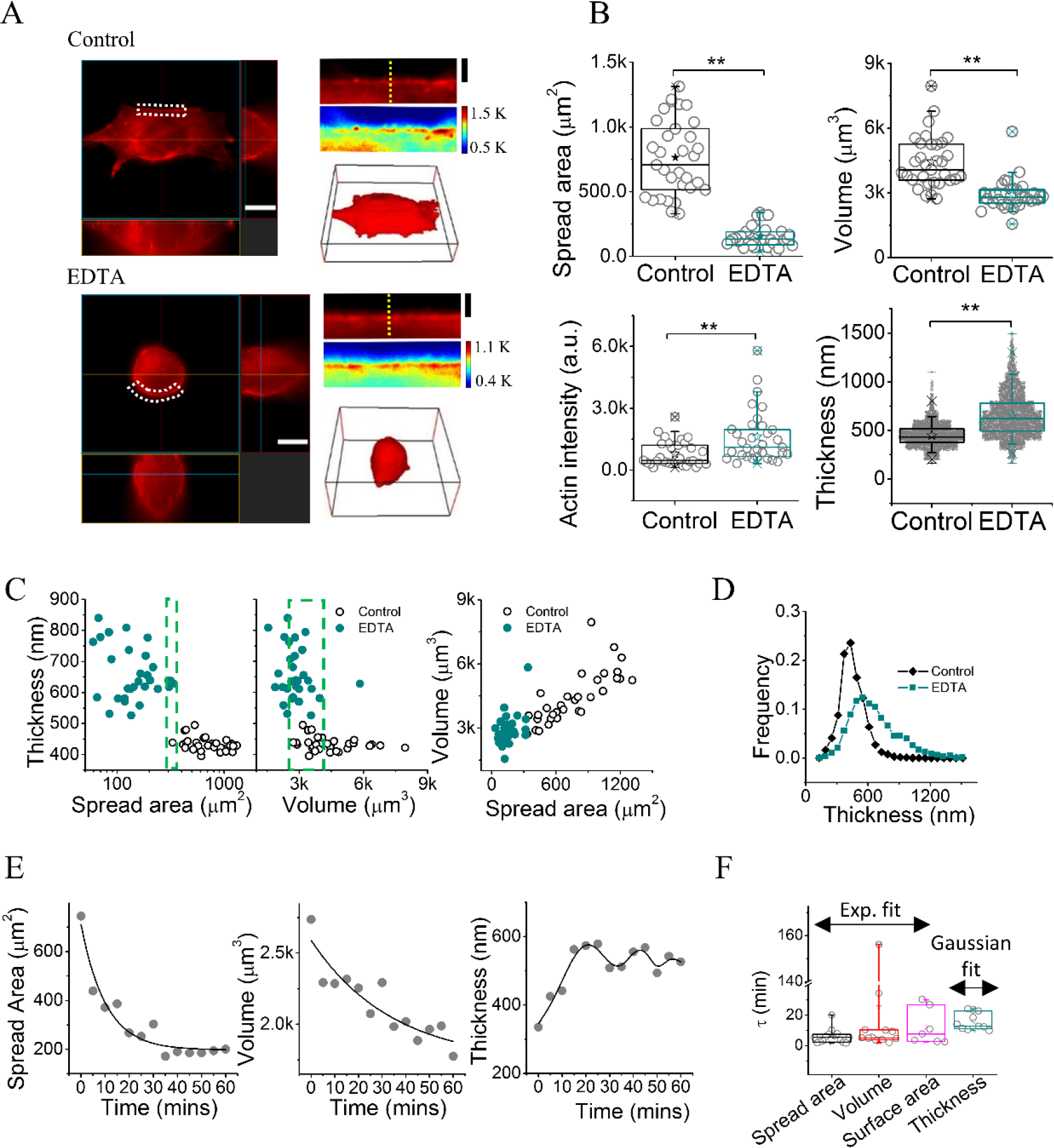
De-adhesion of cell reduces volume and increases cortex thickness. (A) CHO cells transiently expressing Lifeact-mCherry under normal (top) and EDTA-treated (1 hr) conditions. Left: Representative slice view of cells; Right: linearized cortex at white dotted ROI of left panel (top), 3D volume rendered image (bottom). Scale bars, 10 (left) and 2 (right) μm respectively. (B) Spread area, volume, peak actin-intensity and cortex thickness of same LifeAct-mCherry transfected cells before (Control) and after EDTA treatment (n= 32 cells (Control), n=32 cells (EDTA) from three independent experiments). Error bars represent SD. (C) Scatter plot of thickness vs. spread area and thickness (left and centre) and scatter plot of volume vs. spread area for cells represented in (B). (D) Normalized distributions of thickness for both conditions for cells represented in (B). (E) Spread area (left), volume (middle) and thickness (right) of a typical LifeAct-mCherry expressing CHO cell undergoing de-adhesion. Solid lines represent exponential fits to spread area and volume (left, middle) or multi-term Gaussian fit to thickness (right). (F) Average time constants obtained from exponential fits of spread area, volume, surface area evolution and peaking times obtained from Gaussian fits to thickness evolution (n = 14 cells). Wilcoxon based Mann-Whitney U-test were performed for all statistical significant except thickness (one-way ANOVA using Bonferroni post hoc method) with p cut off value < 0.05 and p<0.001 denotes * and **respectively.

For the slower de-adhesion cases (τ∼20 mins), the volume (Fig. 4 E *middle*) reduces at a slower rate (τ∼60-220 mins). However, there are cases where the volume reduction is in the same timescale as the de-adhesion. The cell surface area also decreases with a time constant of ∼5-20 mins (Fig. 4 F). The thickness, however, does not show a monotonous decrease. We find an increase in thickness accompanied by oscillations. The thickness increases with undulations peaking at ∼10-20 mins and 30-40 mins, the peaks identified by multi-term Gaussian fits (Fig. 4 E, *right*). The correlation between volume and thickness (measured every 5 mins through 60 mins) for these cells is higher (∼ −0.15 ± 0.05, averaged over 14 cells) than that in the flat (∼ −0.04) or rounded cells (∼ −0.01).

**FIGURE 5.**
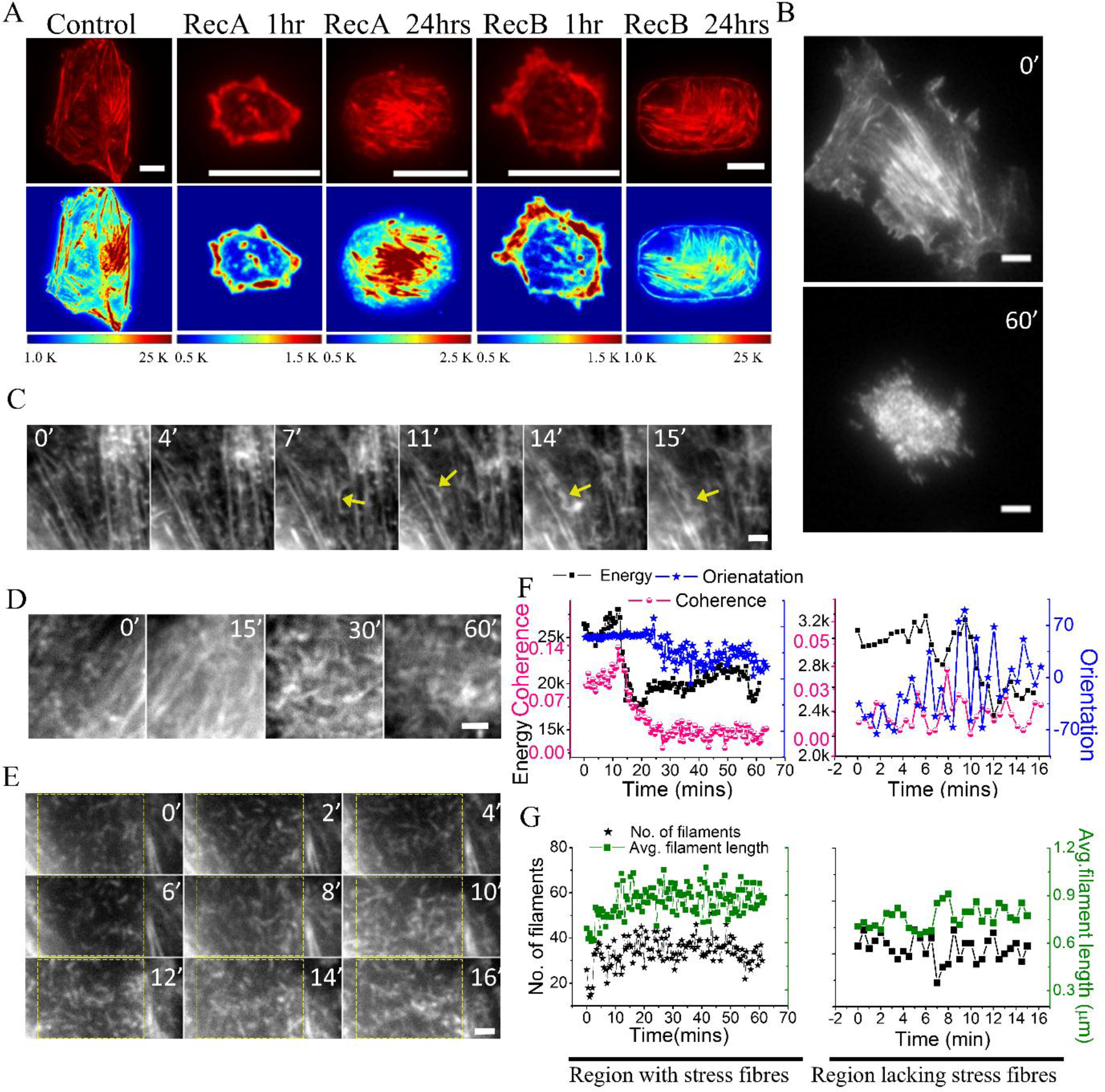
TIRF imaging of basal cortex on spread area alterations. (A) TIRF images of fixed and Rhodamine-phalloidin stained CHO cells grown on glass and micropatterns as indicated. Scale bars, 10 μm. (B) TIRF image of basal cortex of CHO cell transiently expressing LifeAct-mCherry, at indicated. Scale bars, 10 μm. (C) TIRF images showing buckled (marked with yellow arrow) and fragmented of cortical fibres upon de-adhesion. (D) Evolution of a region enriched with stress fibres (E) Evolution of a region lacking stress fibres (F) Evolution of energy (total intensity), coherence and orientation parameters of through de-adhesion in regions depicted in D (left) and yellow ROI of E (right). Scale bars (C-E), 2 μm. Time points (B-G) indicate minutes elapsed after administration of EDTA

### Basal cortex gets fragmented and mobile on de-adhesion

While the measurement of the cortex thickness is carried out at the cell edges (at higher planes than the basal cortex), visualizing the distribution of actin at these regions is difficult. Since visualization of actin may provide important information about processes responsible for thickness increase, we use TIRF microscopy to study the state of cortical-actin at the basal membrane. First, we find that micropatterned cells (Rec A and Rec B) with reduced spread area (as in Fig. 3) lack of stress fibres and have punctuated structures of actin puncta at 1 hr post plating (Fig. 5 A, Fig. S3 A in the Supporting Material). At 24 hrs post plating, with the increase in spread area, there appear more stress fibres in comparison to puncta of actin. Next, we image CHO cells transiently expressing LifeAct-mCherry and find distinct changes on deadhesion (Fig. 5 B-E, Fig. S3 B). Stress fibres vanish in the first 5-15 mins of EDTA treatment (Fig. 5 C) and display curved morphology (Fig. 5 C, yellow arrows), events of buckling (Fig. S4 A in the Supporting Material) and fragmented filaments with increased mobility (Movie S3 in the Supporting Material). We quantify the alterations in actin arrangement by monitoring small regions of cells both with (Fig. 5 D) and without stress fibres (Fig. 5 E) by measuring parameters - orientation, coherence and energy (total intensity) for every frame in the image sequence (29). The frame-to-frame variation of orientation parameter increases after EDTA treatment for both kinds of regions (Fig. 5 F, Fig. S4 B, C). The frame-to-frame variations reflect the change in the average orientation parameter in the 0.5 min time interval between frames. We quantify this by plotting the SD(Orientation) (calculated over 5 consecutive frames) (Fig. S4 D) and find an increase followed by oscillations in most cases implying enhanced dynamics of re-orientation of filaments. In an unperturbed cell that alters its spread area naturally, we find that formation of stress fibres result in the opposite trend - i.e. reduction of frame-to-frame variation in orientation parameter (Fig. S4 E-F). To further characterize the changes in the cortex, we perform object detection (Fig. S4 G) to find out the number and length of filaments. We find a good representation of filaments ranging from 0.3-3 μm (intermediate-sized filaments) at the initial timepoint (Fig. S4 H) and continue analysing their abundance and length. In both types of regions, the number of filaments alter in the first 15 mins on EDTA treatment (Fig. S4 I-J). We either find a biphasic behaviour where the number first decreases (Fig. 5 G) and then increases (6/12 cases, second slope: 1.1 ± 0.75 per min), or, steadily increases (5/12 cases, slope: 0.67 ± 0.45 per min) - although a minor reduction is also observed (1/12 cases, slope: −0.85 per min). While the average filament length increases with time in some cases (Fig. 5 G), it has no particular trend when its evolution for all regions are considered. Regions lacking stress-fibres usually de-adhere and move to central parts of the cell – making long-time behaviour inaccessible. For others, we find that the 25-60 mins interval post EDTA treatment marked a reduction in number of filaments (slope: −0.1 ± 0.06 per min, averaged over 3 cases - 2 reductions and 1 increase) while at 60 mins the number either increases (84 ± 36%) with respect to the beginning or slightly decreases (10%). Together these results suggest that de-adhesion not only induces dissolution (and fragmentation) of stress fibres but causes an increase in the number-density and dynamics of filaments ranging from 0.3-3 μm.

### De-adhesion triggered cortex thickening does not require cell volume reduction

De-adhesion results in stress fibres being contracted internally – suggesting the loss of external pulling forces on the cytoskeleton. Besides the expected reduction in traction forces, reduction in spread area also reduces volume. While spread area decrease may lower the Laplace pressure as reported in liposomes (30) – volume reduction may also further alter the Laplace pressure. It is therefore a matter of investigation if the reduced spread area is sufficient for increasing the thickness or the volume reduction is also essential. To test this out, we first check if volume decrease requires endocytosis. In order to expand the sample-size for measuring volume changes, we adapt an alternative and faster way to assess volume change. We image single cells transiently expressed eGFP or DsRed2, at fixed focal planes, and measure the change in mean intensity of single cells (31) on EDTA treatment. Increased mean intensity implies a higher concentration of the fluorescent protein and hence reports a relative decrease in volume. Although we obtain larger estimates of volume change than z-stack imaging, this technique helps us identify pharmacological agents that hinder the volume reduction on de-adhesion. We observe that volume reduction is hindered by treatments with Cyto D, Jaspla and Blebb (Fig. 6 A-B) that prevent actin polymerization, dynamics (20) and myosin II activity respectively. Dyna treatment – known to block GTPase action of dynamin required for pinching off endocytic buds (21) also hinders volume reduction, implicating endocytosis to be essential for the volume decrease. To validate this further, control and EDTA treated cells are incubated with a membrane dye (FM 1-43 FX) for fixed times and subsequently fixed and imaged to quantify the internal fluorescence (Fig. 6 C). We find that de-adhesion indeed increases the amount of FM internalized – indicating increased endocytosis rate (Fig. 6 D), hence, confirming the role of endocytosis in volume reduction.

**FIGURE 6.**
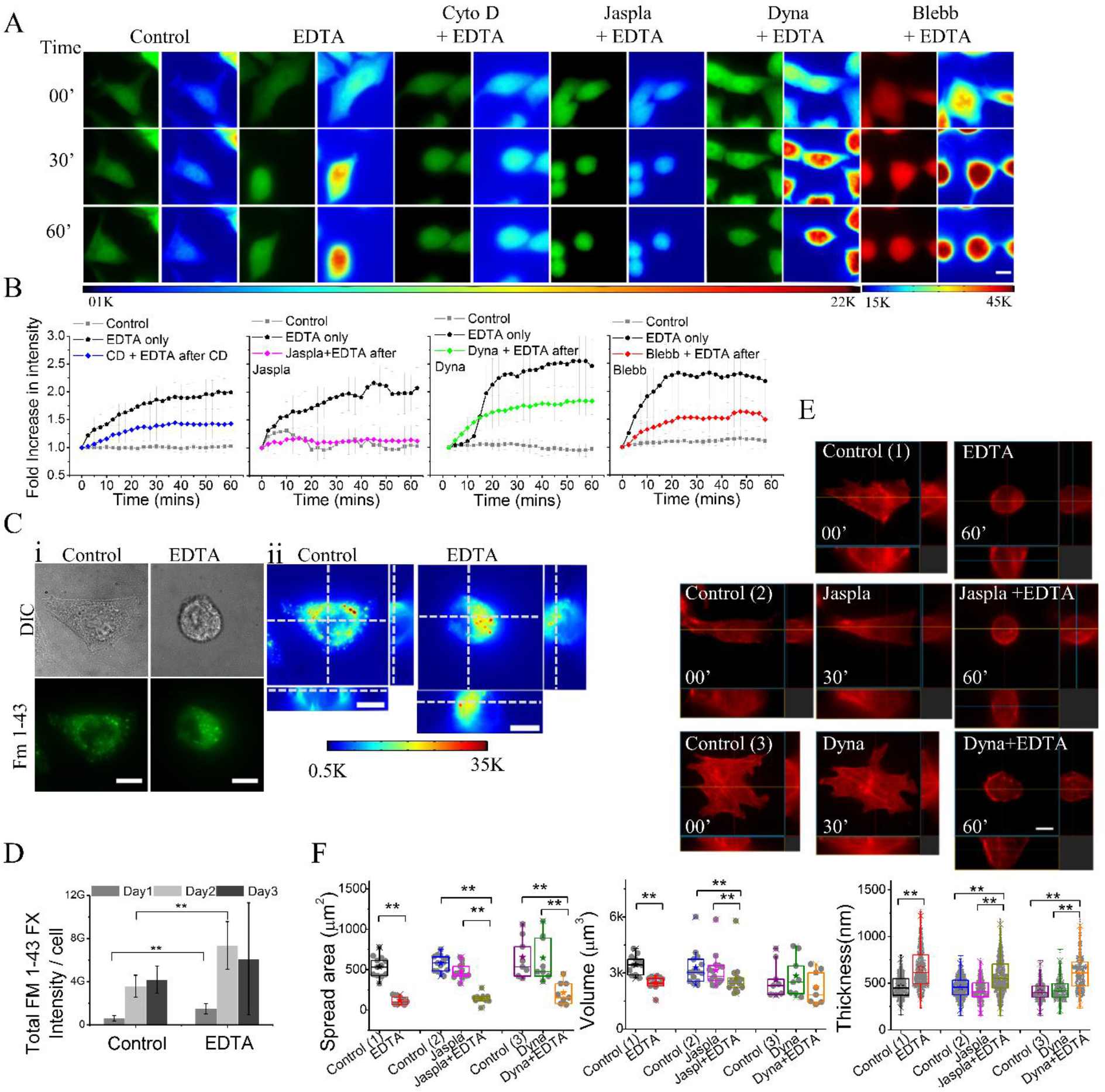
Dependence of de-adhesion induced volume reduction on actomyosin and endocytosis. (A) Time series and corresponding color coded images of CHO cells transiently expressing eGFP or DsRed2 (right-most only) in different conditions and time points as indicated. Scale bar, 10 μm. (B) Evolution of mean cytosolic intensity in Control, EDTA and Cyto D + EDTA (average of n= 15, 13 and 14 cells respectively from three independent experiments); Control, EDTA and Jaspla + EDTA (average of n= 03, 04 and 14 cells respectively from one experiment) and Control, EDTA and Dyna + EDTA (average of n= 13, 6 and 21 cells respectively form three independent experiments); and Control, EDTA and Blebb + EDTA (n= 05, 11 and 16 cells respectively from two independent experiments) conditions. (C) (i) DIC (top) and epifluorescence (bottom) images of CHO cells labelled with FM 1-43 FX dye for 1hr in control conditions and after 1 hr of EDTA treatment with their (ii) slice views (xy, xz and yz planes). Scale bar, 10 μm. (D) Total FM 1-43 FX intensity per cell in different conditions on different days (Day1 n= 32 and 31, Day2 n= 20 and 21 and Day3 n= 15 and 13 cells for Control and EDTA) (p<0.001). (E) Slice view of LifeAct-mCherry transfected CHO cells on three different conditions as indicated. Scale bars, 10 μm. (F) Spread area (left), volume (middle) and thickness (right) of cells in the indicated conditions. Wilcoxon based Mann-Whitney U-test were performed for all statistical significant except thickness (oneway ANOVA using Bonferroni post hoc method) with p cut off value < 0.05 and p<0.001 denotes * and **respectively.

In these conditions where de-adhesion results in a decrease of spread area but the volume change is reduced, we next study the effect on the cortex thickness by imaging same cells before and after de-adhesion. Only Jaspla and Dyna (Fig. 6 E) are chosen so that the cortex structure is not completely destroyed and the thickness is also not affected by the drugs before de-adhesion. Although spread area decreases (Fig. 6 F *left*), we find that the volume change on EDTA treatment (in the continuous presence of Jaspla and Dyna) decreases from 28% for control to 14% for Jaspla treated and 21% for Dyna treated. Dyna alone results in increased volume (8%) as clear in Fig. 6 F (*middle*) (Table S1 in the Supporting Material). To understand the combined effect of Dyna and EDTA, the volume altered is computed with respect to the control condition (absence of Dyna). The net volume decreases due to combined effect of Dyna and EDTA is only 15% - closely matching the effect of Jaspla (18% reduction when Jaspla + EDTA is compared with Control condition). Therefore, following single cells confirms that Dyna and Jaspla hinder volume reduction on deadhesion.

Cortex thickness, however, is enhanced on EDTA treatment for untreated (42%), Jaspla treated (36%) as well as Dyna treated cells (41%) (Fig. 6 F *right*, Table S1). These results imply that cortex thickening on de-adhesion does not require a concomitant reduction in volume. However, we also find that cortical volume – estimated from the product of cortex thickness and cell’s surface area - decreases significantly by 10% for control cells after 1 hr of EDTA treatment. Although thickness alters significantly, the cortical volume shows no significant change under the continued presence of Dyna and Jaspla treated cells. Volume reduction hence is not crucial for cortex thickening but important for further modification of the cortex - indicating existence of later phases of cortex remodeling.

Together, the results emphasize that cortex responds to reduction of spread area – going through remodeling marked by dissolution of stress fibres; increased dynamics and greater abundance of intermediate-sized fibres; oscillations in thickness; and alteration of the cortical volume. We demonstrate that transiently increased contractility decreases thickness and *vice versa* while triggering lateral contraction increases the thickness and results in oscillations.

## Discussion

The thin layer of actin network underlying the plasma membrane is found, to the best of our knowledge, in all nucleated cells of different size and shapes. In this work, we study cortical thickness regulation in well-spread CHO cells by perturbing actomyosin activity and restricting cell’s spread area (Table S2 in the Supporting Material).

The tight regulation of cortex thickness is reflected in its low intracellular and intercellular variability (SD/mean ∼ 0.15) and its increase when regulatory factors are perturbed. We show that treatments that reduce actin polymerization (Cyto D, Lat B) finally result in increased thickness. We believe the treatments weaken the cortex (8, 32), lead to transient build-up of tension due to increased contractility, as expected from higher myosin II to F-actin ratio (33, 34) and evident from the straightening of the cell-edge (Fig. S2 F). Transiently decreased cortex thickness (Fig. S2 G-H) points to the inverse correlation between contractility and thickness. We find that this phase is followed by cortex rupture and heterogeneous lateral contraction of the cortex (Movies S1 and S2) with increased average cortex thickness and variability. The inverse relation between thickness and contractility, is reinforced by observations of increased thickness on Blebb treatment. The increased intensity on ATP depletion/Blebb treatment further imply active regulation of F-actin concentration at the cortex. This is consistent with studies reporting increased F-actin population on ATP depletion (35, 36) and also in line with the role of myosin II in actin disassembly sites in keratocytes (37).

We have further reported that spread area alters cortex thickness. Cells on micropatterns with smaller adhesive area have thicker cortex showing that well-spread interphase cells have a thinner cortex than when rounded. Restricting cells in small micropatterns also resulted in a significant volume decrease which is also observed when cells are de-adhered – in line with past reports (38). We show that this volume reduction is due to endocytosis and requires actomyosin dynamics - as expected from their role in endocytosis (15). The volume reduction however doesn’t crucially control cortex thickening on de-adheison. When we highlight (green box, Fig. 4 C) the parameter range in which cells have the same spread area/volume but different thickness, we find the overlap is smaller for spread area than for volume implying spread area has a larger and more critical role in thickness determination than volume. We also evidence cortex thickening despite hindering volume reduction (using Jaspla/Dyna). Together, these results indicate that the process of de-adhesion itself may have the necessary cues for the cortex thickening. We, therefore, believe that the loss of surface forces may trigger the initial remodeling of cortex.

The remodeling of the cortex persists beyond the initial thickening and is reflected in the reduced cortical volume on EDTA treatment on control but not Jaspla/Dyna treated cells. Oscillations in thickness following the initial increase (during de-adhesion of control cells) and biphasic (or even oscillating) trends of filament number and length in single filament analysis of basal cortex also confirm the existence of later stages of remodeling.

The remodeling of the actin network organization by micropatterning/de-adhesion is studied by TIRF imaging of the basal cortex. Stress fibres – observed in majority of CHO cells, are absent in cells nascently adhered to micropatterns (Fig. 5 A) and also disintegrated by de-adhesion. We believe this is caused by fragmentation of stress fibres because we evidence buckling of stress fibres and find increased number of both dynamic and distorted filaments (Movie S3) on de-adhesion. Regions lacking stress fibres also display enhanced dynamics and a growing population of intermediate-sized fibres which are however, uncorrelated to each other (Fig. 5 F r, G *right*, Fig. S5 B, C *bottom*). This implies that de-adhesion increases dynamics of the existing filaments while fragmenting and creating more filaments. These results help us postulate that the sudden loss of pulling forces by the substrate (at the end of stress fibers) allows myosin-II-based contractility to take over – leading to larger inward forces and hence buckling and fragmenting the cortical network. This response is also expected to result in lateral contraction of the cortex as well as increased dynamics of single filaments – both of which are observed. Similar unhindered lateral contraction is also observed just after rupture of the Cyto D treated cell cortex.

Together, this implies that increased contractility decreases thickness of the cortex only when lateral contraction is not possible due to the ends being rigidly fixed. The resulting build-up of tension is expected to first straighten and align filaments parallel to themselves and thus reduce thickness. When ends are no more fixed, lateral contraction is triggered which may increase events of filament -buckling/fragmenting (39), causing their misalignment and hence increasing cortex thickness.

However, we need to understand if the accumulation of filaments in a smaller surface area due to lateral contraction, is the main cause of cortex thickening. If this were true, the net cortical volume would have still been constant after de-adhesion - which we show (Table S1) is not the case. Cortical volume is reduced actively after EDTA treatment indicating that other processes deplete the accumulated cortex. We believe, actomyosin turnover remodels the thickened cortex and the final thickness depends on the evolving force balance. The persistence of thickened cortex even at 24 hrs post plating on micropatterns and thickness oscillations (Fig. 4 E) at 10-60 mins after de-adhesion confirm this.

Therefore, in this work, we demonstrate that spread area and contractility together regulate cortex thickness and postulate that this is due to the altered impact of myosin-II-based contractility on the actin network. Thus, well-spread interphase cells present a particular organization of the cortex – different from rounded cells. The strong interaction of their cortex with the ECM results not only in an inverse relation of thickness and contractility but also makes their cortex thickness sensitive to spread area.

## Acknowledgements

We thank Jayasri Das Sarma for CHO cells and Arikta Biswas for critical reading of the manuscript. We thank Arikta Biswas and Darius Köster for discussions. RK thanks UGC for his doctoral fellowship. This work was supported by the Wellcome Trust/DBT India Alliance fellowship (grant number IA/I/13/1/500885) awarded to BS.

## Author Contribution

BS conceptualized the project. BS and RK set up the analysis. RK performed and analysed the experiments. BS and RK interpreted the data and wrote the paper.

## Supporting material

**Figure S1.**
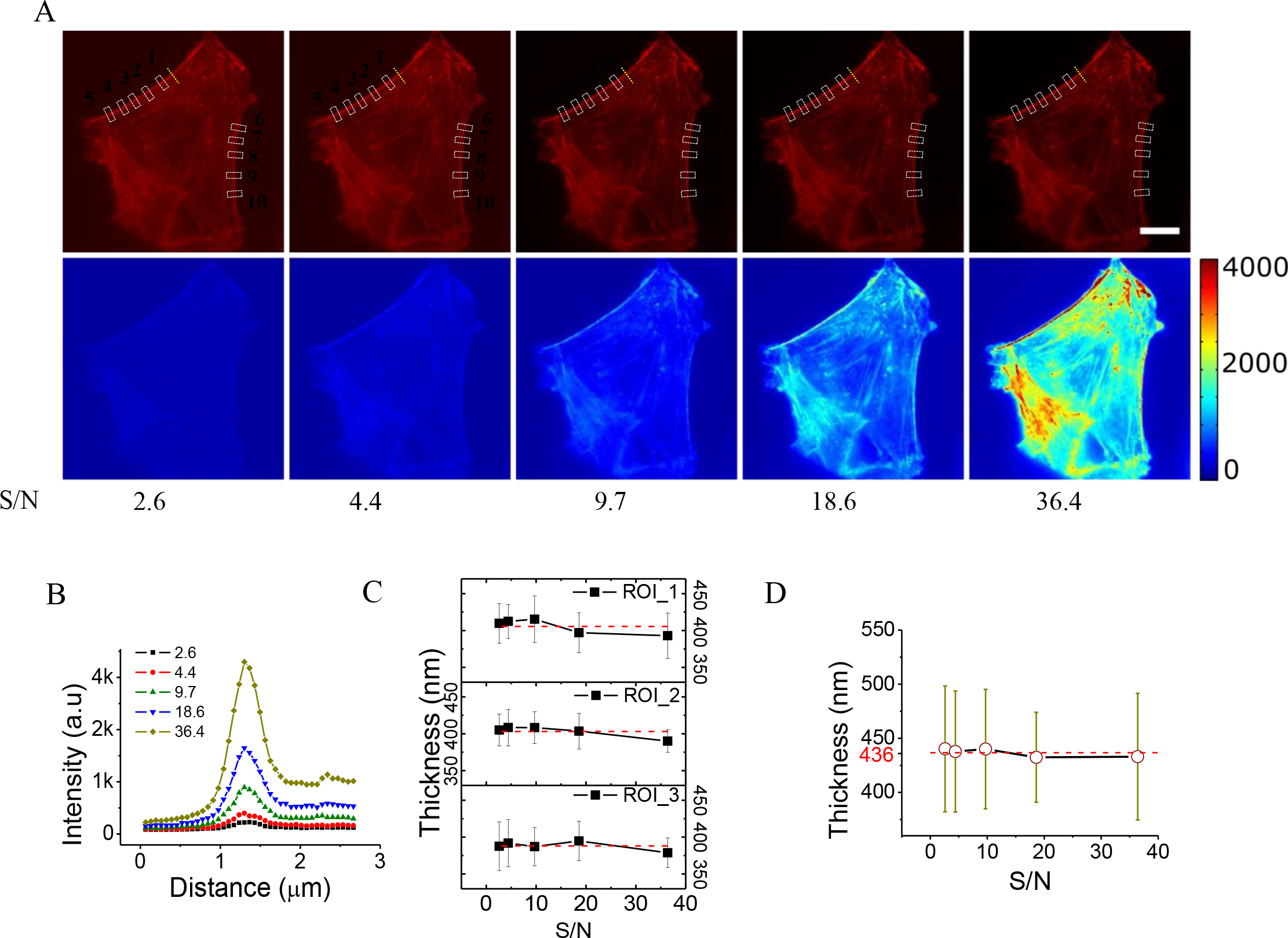
Dependence of thickness measurement on signal-to-noise (S/N) of images. (A) Rhodamine-phalloidin stained image of a CHO cell captured at different exposure times (top) and corresponding color coded image (bottom) with the S/N mentioned below. Scale bar, 10 μm. (B) Intensity line-scan of F-actin normal to the cortex for the same region but different S/N. (C) Plot of average thickness change on ROIs 1, 2 and 3 over S/N (as depicted in dotted rectangle in (A)). (D) Dependence of average thickness (of 10 ROIs) on the S/N at which images were captured. (Error bars represent SD).

**Figure S2.**
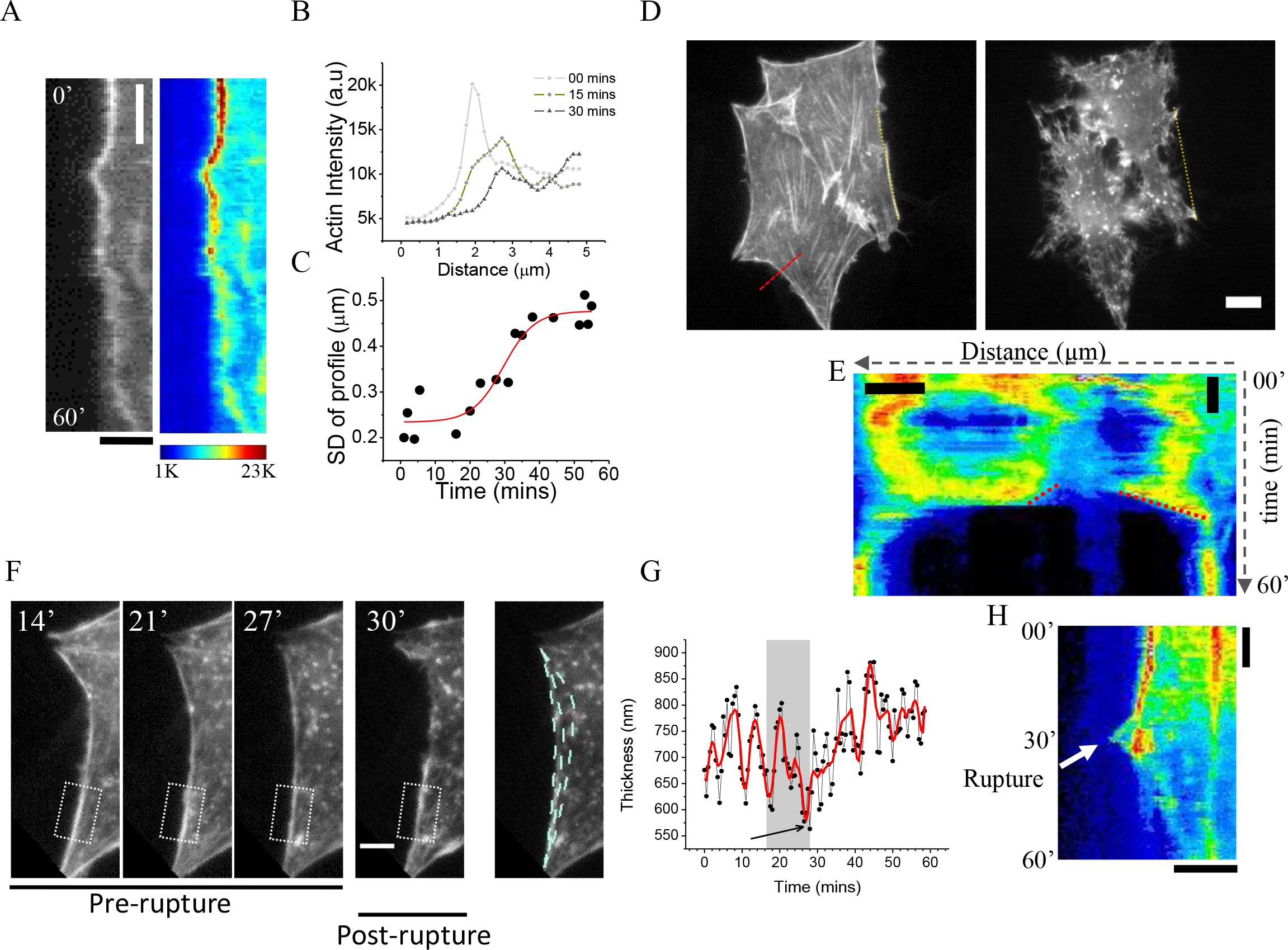
Evolution of cortex thickness on Cyto D treatment. (A) Kymograph of a line normal to the cortex of LifeAct-mCherry transfected CHO cells (Movie S2) after Cyto D administration at time t=0 min. Scale bars, y=10 mins and x=3 μm. (B) Representation of intensity profile across the line represented in (A) at three time points after Cyto D addition. (C) The evolution of cortex thickness across the line represented in (A). Line denotes sigmoidal fit. (D) CHO cells transiently expressing LifeAct-mCherry before (left) and after 60 mins (right) of Cyto D treatment (Movie S1). Scale bar, 10 μm. (E) Kymograph of cell in (D) at ROI overlaid in yellow. Red lines mark the movement of ends of ruptured cortex, used to estimate the velocity of contraction. Scale bars, x = 5 μm and y=10 mins. (F) Image sequence showing a small curved region of cortex straightening, finally rupturing. White dotted line overlaid highlights the contours of the edge while straightening (*right*). (G) Average thickness of a region of the cortex (shown in dotted white box (F)) followed over time. Shaded region highlights time-period at which curved region straightens just prior to its rupture. (H) Kymograph of cells in (D) at ROI overlaid in red line. Arrow showing points at which cortex ruptures.

**Figure S3.**
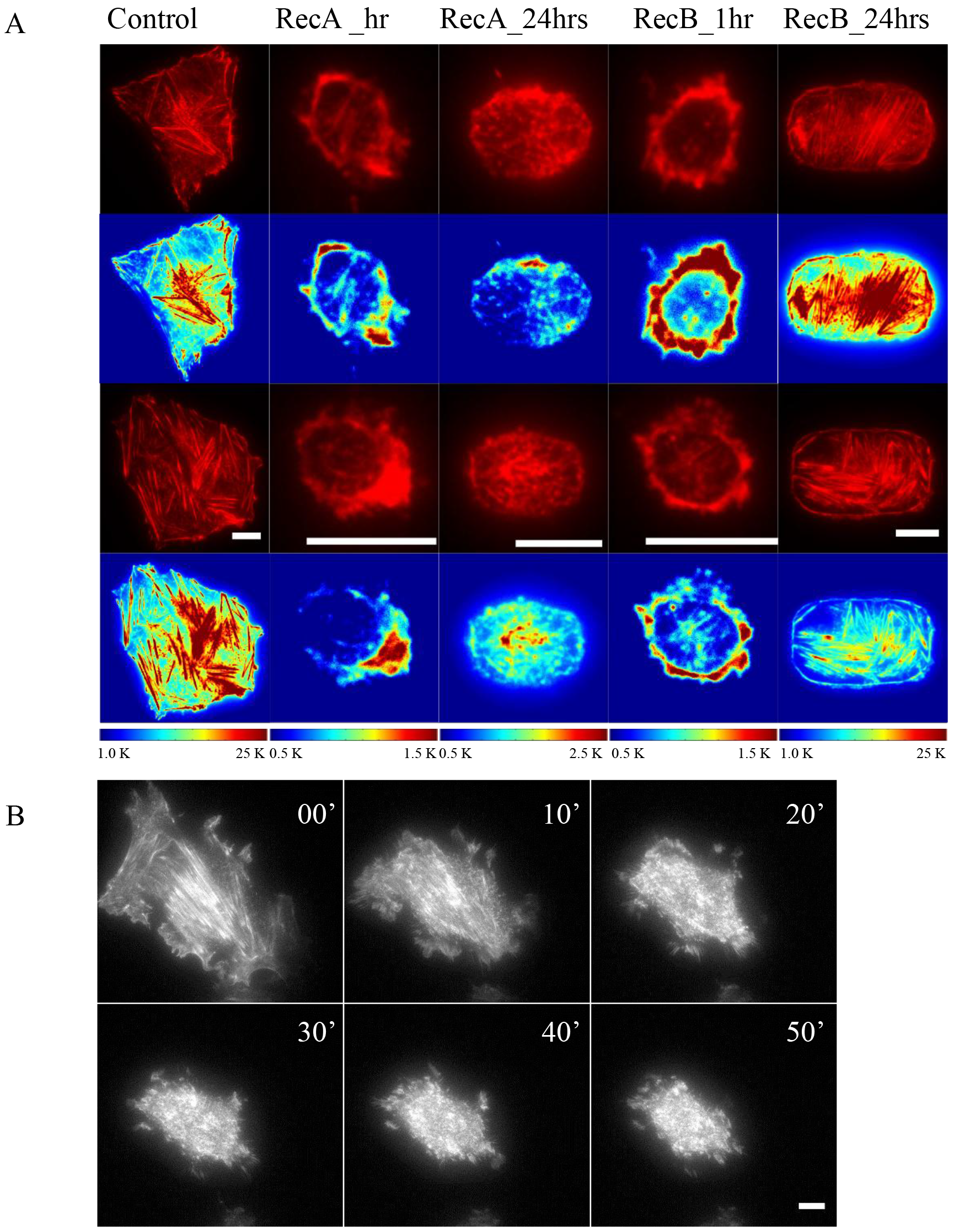
TIRF images of cortex with altered spread area. (A) TIRF microscopy images of Rhodamine-phalloidin stained CHO cells (*top*) grown on different micropatterns and time-points post plating as indicated with their corresponding color coded images (*bottom*). (B) Time lapse TIRF images of basal cortex of CHO cells expressing LifeAc-mCherry during de-adhesion. Scale bars,10 μm.

**FIGURE S4.**
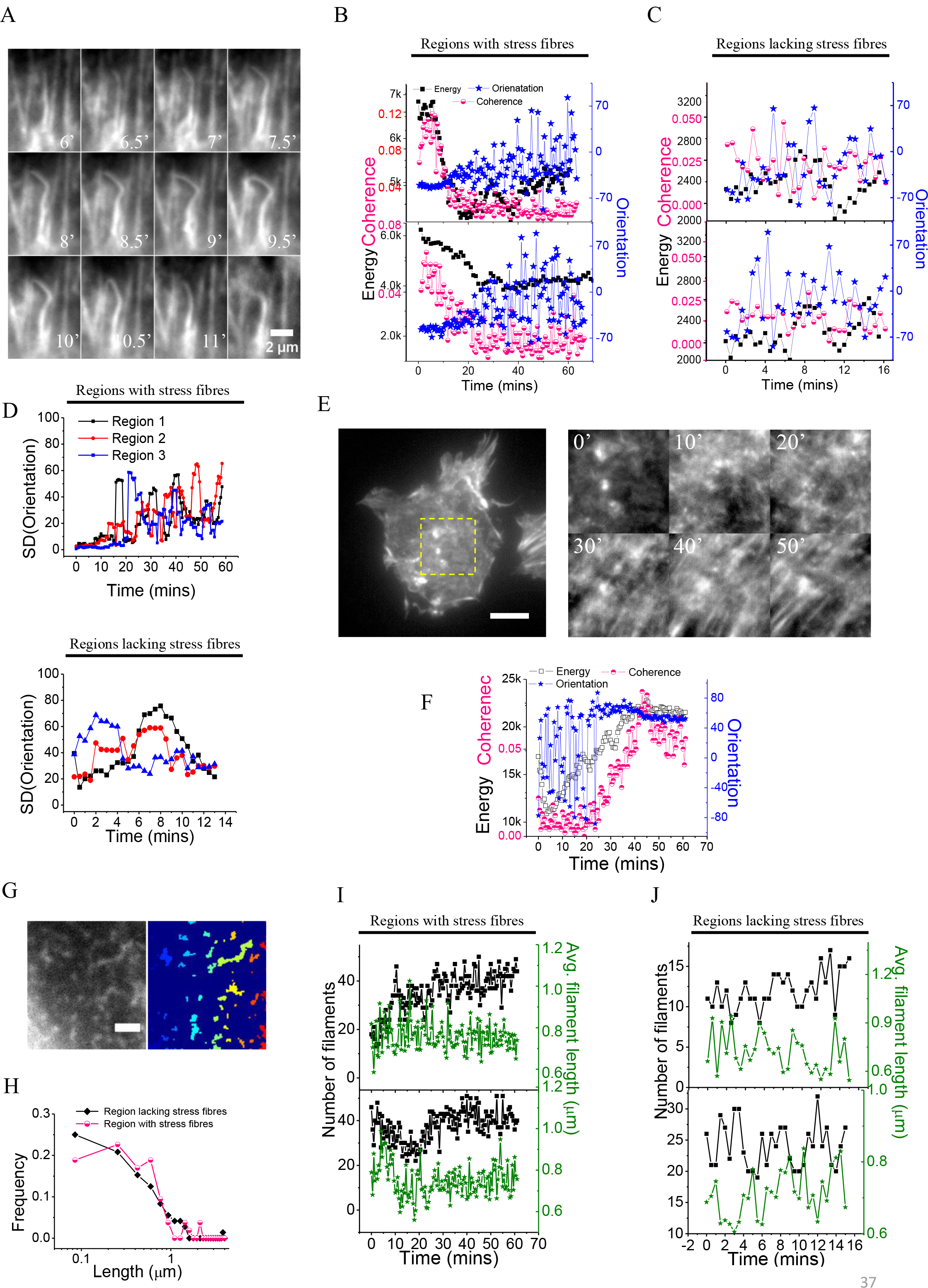
TIRF imaging and quantification of actin re-arrangement in CHO cells expressing LifeAct-mCherry. (A) Zoomed view showing buckling event of single fibre during de-adhesion (EDTA administered at t=0). Scale bar, 2 μm. (B,C) Evolution of energy, orientation and coherence upon de-adhesion measured in regions with stress fibres (B) and regions lacking (C) them (EDTA administered at t=0). (D) Evolution of SD(orientation) at regions with (*top*) and lacking (*bottom*) stress fibres. (E) *Left*: Cell altering spread area naturally. Scale,10 μm. *Right*: Temporal evolution of region in yellow in the *left* image. (F) Evolution of energy, orientation and coherence of images depicted in E, *right*. (G) Representative gray-scale (*left*) and object-labelled (right) images where single filaments are detected and represented in different colours. Scale bar, 2 μm. (H) Normalized distribution of filament size measured in region with and lacking stress fibres. (I, J) Evolution of number and average length of intermediate-sized filaments for two region with (I) and lacking stress (J) fibres during cell deadhesion (EDTA administered at t=0).

**Table S1.** Comparison of cell and cortex parameters for different conditions. The cell volume, sphericity, surface area, spread area and cortex thickness and cortical volume for different conditions are computed for single cells followed before and after (treatment and) EDTA and then averaged. For the estimation of the fraction of cortical volume left after EDTA treatment, the ratio of (surface area multiply by thickness) is computed for EDTA treated cells with respect to control. When control was the untreated cell no colours highlight the box for hypothesis testing. When the control is the drug treated cell, the box is highlighted in pink or green depending on the drug treatment. One sample t-test with test mean =1 and alternate mean <>1 was used to obtain the *p* values. NS signifies *p* values above 0.05.

| | Condition | Average Thickness (nm)<br>(*h*) | Average Volume (μm³) | Average Spread area (μm²) | Average Surface area<br>(*A*) | Average Sphericity | Fraction of cortical volume left<br>$= \frac{(A * h)_{EDTA}}{(A * h)_{Control}}$<br>(*Hypothesis testing*) | | No. of Cells followed |
| --- | --- | --- | --- | --- | --- | --- | --- | --- | --- |
| 1 | Control (1) | 466 | 3459 | 536 | 1443 | 0.718 |  |  | 12 |
|  | EDTA | 662 | 2488 | 120 | 903 | 0.918 | 0.90<br>(*p*=0.0355) |  |  |
| 2 | Control (2) | 455 | 3291 | 579 | 1457 | 0.674 |  |  | 14 |
|  | Jaspla | 427 | 3146 | 475 | 1332 | 0.718 |  |  |  |
|  | Jaspla + EDTA | 584 | 2692 | 137 | 1011 | 0.890 | 0.95<br>(NS) | 1.06<br>(NS) |  |
| 3 | Control (3) | 415 | 2642 | 665 | 1339 | 0.646 |  |  | 8 |
|  | Dyna | 434 | 2855 | 605 | 1337 | 0.714 |  |  |  |
|  | Dyna + EDTA | 615 | 2249 | 215 | 924 | 0.842 | 1.03<br>(NS) | 1.01<br>(NS) |  |

**Table S2.** Details of statistical parameters. Table provide details of the statistical parameter for data presented in main and supplementary figures. Two methods were used for finding significance difference. * Represent one-way ANOVA using Bonferroni post hoc method and # for unpaired Wilcoxon based Mann-Whitney U test. Here, N1(Cells) represent total number of cells including all days were used in analysis and N1 (ROIs) signifies the number of ROIs (length of each ROIs was 2 μm) for each cell were considered.

| Figure 1 D |  |  |  |  |  |  |  |  |
| --- | --- | --- | --- | --- | --- | --- | --- | --- |
| Parameter |  | N1(cells) | N2(ROIs) | Mean (nm) | Median | SD | SEM | p-value (wrt control) |
| Thickness |  | 1 | 10 | 381 | 381 | 57 |  |  |
| Thickness |  | 30 | 10 | 424 | 417 | 62 |  |  |
| Figure 2 D |  |  |  |  |  |  |  |  |
| Parameter | Condition | N1(cells) | N2(ROIs) | Mean (a.u) | Median (a.u) | SD (a.u) | SEM (a.u) | p-value (wrt control) |
| Actin intensity | Control | 30 | 10 | 21966 | 21466 | 6975 | 70 |  |
|  | Cyto D | 30 | 10 | 9110 | 7866 | 5136 | 52 | *0 |
|  | Lat B | 30 | 10 | 13378 | 12632 | 6592 | 66 | *0 |
|  | ATP dep | 30 | 10 | 32643 | 30000 | 13904 | 141 | *0 |
|  | Blebb | 30 | 10 | 29570 | 28683 | 9070 | 91 | *0 |
| Figure 2 E |  |  |  |  |  |  |  |  |
| Parameter | Condition | N1(cells) | N2(ROIs) | Mean (nm) | Median (nm) | SD (nm) | SEM (nm) | p-value (wrt control) |
| Thickness | Control | 30 | 10 | 425 | 401 | 108 | 2 |  |
|  | Cyto D | 30 | 10 | 483 | 522 | 182 | 3 | *3.8x10<sup>-127</sup> |
|  | Lat B | 30 | 10 | 541 | 461 | 226 | 4 | *4.7x10<sup>-302</sup> |
|  | ATP dep | 30 | 10 | 557 | 521 | 190 | 3 | *6.5x10<sup>-187</sup> |
|  | Blebb | 30 | 10 | 515 | 476 | 175 | 3 | *3.4x10<sup>-90</sup> |
| Figure 3 B |  |  |  |  |  |  |  |  |
| Parameter | Condition | N1(cells) |  | Mean (μm<sup>2</sup>) | Median (μm<sup>2</sup>) | SD (μm<sup>2</sup>) | SEM (μm<sup>2</sup>) | p-value (wrt control) |
| Spread area (μm<sup>2</sup>) | Control | 40 |  | 957 | 990 | 223 | 35 |  |
|  | RecA_1h | 40 |  | 83 | 78 | 39 | 6 | #1.4x10<sup>-14</sup> |
|  | RecA_24h | 40 |  | 193 | 201 | 56 | 9 | #1.4x10<sup>-14</sup> |
|  | RecB_1h | 40 |  | 65 | 67 | 21 | 3 | #1.4x10<sup>-14</sup> |
|  | RecB_24h | 40 |  | 432 | 441 | 85 | 14 | #2.3x10<sup>-14</sup> |
| Figure 3 D |  |  |  |  |  |  |  |  |

| Parameter | Condition | N1(cells) |  | Mean (μm<sup>3</sup>) | Median (μm<sup>3</sup>) | SD (μm<sup>3</sup>) | SEM (μm<sup>3</sup>) | p-value (wrt control) |
| --- | --- | --- | --- | --- | --- | --- | --- | --- |
| Volume (μm<sup>3</sup>) | Control | 40 |  | 4873 | 4538 | 1241 | 196 |  |
|  | RecA_1h | 40 |  | 1774 | 1709 | 440 | 70 | #1.4x10<sup>-14</sup> |
|  | RecA_24h | 40 |  | 2493 | 2414 | 552 | 87 | #6.8x10<sup>-14</sup> |
|  | RecB_1h | 40 |  | 1964 | 1937 | 396 | 63 | #1.4x10<sup>-14</sup> |
|  | RecB_24h | 40 |  | 3242 | 2910 | 945 | 149 | #3.0x10<sup>-8</sup> |
| Figure 3 E |  |  |  |  |  |  |  |  |
| Parameter | Condition | N1(cells) | N2(ROIs) | Mean (nm) | Median (nm) | SD (nm) | SEM (nm) | p-value (wrt control) |
| Thickness | Control | 40 | 10 | 430 | 409 | 111 | 2 |  |
|  | RecA_1h | 40 | 10 | 592 | 548 | 204 | 3 | *0 |
|  | RecA_24h | 40 | 10 | 536 | 499 | 179 | 3 | 4.4x10<sup>-168</sup> |
|  | RecB_1h | 40 | 10 | 592 | 554 | 202 | 3 | *0 |
|  | RecB_24h | 40 | 10 | 507 | 474 | 160 | 3 | 1.4x10<sup>-89</sup> |
| Figure 4 B |  |  |  |  |  |  |  |  |
| Parameter | Condition | N1(cells) |  | Mean (μm<sup>2</sup>) | Median (μm<sup>2</sup>) | SD (μm<sup>2</sup>) | SEM (μm<sup>2</sup>) | p-value (wrt control) |
| Spread area (μm<sup>2</sup>) | Control | 32 |  | 764 | 741 | 291 | 51 |  |
|  | EDTA | 32 |  | 155 | 134 | 81 | 14 | #7.2x10<sup>-12</sup> |
| Parameter | Condition | N1(cells) |  | Mean (μm<sup>3</sup>) | Median (μm<sup>3</sup>) | SD (μm<sup>3</sup>) | SEM (μm<sup>3</sup>) | p-value (wrt control) |
| Volume (μm<sup>3</sup>) | Control | 32 |  | 4432 | 4188 | 1202 | 212 |  |
|  | EDTA | 32 |  | 2914 | 2814 | 709 | 125 | #2.2x10<sup>-8</sup> |
| Parameter | Condition | N1(cells) | N2(ROIs) | Mean (a.u) | Median (a.u) | SD (a.u) | SEM (a.u) | p-value (wrt control) |
| Actin intensity | Control | 32 | 10 | 786 | 868 | 606 | 107 |  |
|  | EDTA | 32 | 10 | 1606 | 1148 | 1273 | 225 | #8.5x10<sup>-4</sup> |

**1 Table S2. Details of statistical parameters.**
| Parameter | Condition | N1(cells) | N2(ROIs) | Mean (nm) | Median (nm) | SD (nm) | SEM (nm) | p-value (wrt control) |
| --- | --- | --- | --- | --- | --- | --- | --- | --- |
| Thickness | Control | 32 | 10 | 448 | 432 | 115 | 2 |  |
|  | EDTA | 32 | 10 | 657 | 619 | 224 | 4 | *3.3x10<sup>-16</sup> |
| Figure 6 D |  |  |  |  |  |  |  |  |
| Parameter | Condition | N1(cells) |  | Mean (a.u) | Median (a.u) | SD (a.u) | SEM (a.u) | p-value (wrt control) |
| Total FM1-43 FX intensity /cell_Day1 | Control | 33 |  | 6.0x10<sup>8</sup> | 5.0x10<sup>8</sup> | 2.3x10<sup>8</sup> | 4.1x10<sup>6</sup> |  |
|  | EDTA | 32 |  | 1.5x10<sup>9</sup> | 1.5x10<sup>9</sup> | 4.7x10<sup>9</sup> | 8.5x10<sup>8</sup> | *9.8x10<sup>-14</sup> |
| Total FM1-43 FX intensity /cell_Day2 | Control | 22 |  | 3.5x10<sup>9</sup> | 3.4 x10<sup>9</sup> | 3.5x10<sup>9</sup> | 2.2 x10<sup>8</sup> |  |
|  | EDTA | 21 |  | 7.3x10<sup>9</sup> | 6.7 x10<sup>9</sup> | 7.3 x10<sup>9</sup> | 4.8 x10<sup>9</sup> | *3.9x10<sup>-8</sup> |
| Total FM1-43 intensity /cell_Day3 | Control | 16 |  | 4.1 x10<sup>9</sup> | 4.1 x10<sup>9</sup> | 4.1 x10<sup>9</sup> | 3.2 x10<sup>8</sup> |  |
|  | EDTA | 14 |  | 6.14x10<sup>9</sup> | 4.3 x10<sup>9</sup> | 6.1 x10<sup>9</sup> | 1.4x10<sup>8</sup> | *1.6x10<sup>-1</sup> |
| Figure 6 F |  |  |  |  |  |  |  |  |
| Parameter | Condition | N1(cells) |  | Mean (µm<sup>2</sup>) | Median (µm<sup>2</sup>) | SD (µm<sup>2</sup>) | SEM (µm<sup>2</sup>) | p-value (wrt control) |
| Spread area (µm<sup>2</sup>) | Control (1) | 12 |  | 537 | 541 | 130 | 38 |  |
|  | EDTA | 12 |  | 120 | 111 | 49 | 14 | #3.7x10<sup>-5</sup> |
|  | Control (2) | 14 |  | 537 | 585 | 130 | 38 |  |
|  | Jaspla | 14 |  | 475 | 449 | 91 | 24 |  |
|  | Jaspla + EDTA | 14 |  | 136 | 144 | 60 | 16 | #7.4x10<sup>-6</sup> |
|  | Control (3) | 8 |  | 537 | 572 | 130 | 38 |  |
|  | Dyna | 8 |  | 646 | 507 | 279 | 99 |  |
|  | Dyna+EDTA | 8 |  | 215 | 173 | 137 | 49 | #1.0x10<sup>-3</sup> |
| Parameter | Condition | N1(cells) |  | Mean (µm<sup>3</sup>) | Median (µm<sup>3</sup>) | SD (µm<sup>3</sup>) | SEM (µm<sup>3</sup>) | p-value (wrt control) |
| Volume<br>($\mu\text{m}^3$) | Control (1) | 12 | | 3460 | 3445 | 492 | 142 | |
| | EDTA | 12 | | 2489 | 2516 | 354 | 102 | #$4.8 \times 10^{-4}$ |
|  | Control (2) | 14 |  | 3291 | 3065 | 952 | 254 |  |
|  | Jaspla | 14 |  | 3147 | 2964 | 914 | 244 |  |
| | Jaspla +<br>EDTA | 14 | | 2693 | 2484 | 954 | 255 | #$1.2 \times 10^{-4}$ |
|  | Control (3) | 8 |  | 2642 | 2364 | 949 | 335 |  |
|  | Dyna | 8 |  | 2855 | 2565 | 1082 | 383 |  |
| | Dyna+EDTA | 8 | | 2249 | 2020 | 873 | 308 | #$0.7 \times 10^{-3}$ |
| **Parameter** | **Condition** | **N1(cells)** | **N2(ROIs)** | **Mean**<br>**(nm)** | **Median**<br>**(nm)** | **SD**<br>**(nm)** | **SEM**<br>**(nm)** | **p-value**<br>**(wrt**<br>**control)** |
| Thickness | Control (1) | 12 | 10 | 466 | 448 | 120 | 4 |  |
| | EDTA | 12 | 10 | 662 | 614 | 212 | 6 | *$4.9 \times 10^{-4}$ |
|  | Control (2) | 14 | 10 | 455 | 455 | 115 | 4 |  |
|  | Jaspla | 14 | 10 | 427 | 408 | 117 | 4 |  |
| | Jaspla +<br>EDTA | 14 | 10 | 584 | 551 | 195 | 5 | *$1.2 \times 10^{-4}$ |
|  | Control (3) | 8 | 10 | 415 | 398 | 104 | 4 |  |
|  | Dyna | 8 | 10 | 434 | 416 | 118 | 5 |  |
| | Dyna+EDTA | 8 | 10 | 615 | 610 | 187 | 6 | *$1.0 \times 10^{-4}$ |

**Movie S1: Effect of Cyto D on actin cortex imaged at 2 frames per minute**.

The movie shows the effect of Cyto D (10 μM) on CHO cells transiently expressing LifeAct-mCherry. Cyto D is administered at t=0.

**Movie S2: Effect of Cyto D on actin cortex imaged at 1 frame per minute**.

The movie shows the effect of Cyto D (10 μM) on CHO cells transiently expressing LifeAct-mCherry. Cyto D is administered at t=0.

**Movie S3: Effect of deadhesion on actin cortex imaged in TIRF at 2 frames per minute**.

The movie shows the effect of EDTA (1 mM) on CHO cells transiently expressing LifeAct-mCherry. EDTA is administered at t=0.

